# GABAergic TH2 Amacrine Cells Participate in Spontaneous Wave Activity in the Developing Retina

**DOI:** 10.64898/2026.02.06.704451

**Authors:** Wenqiu Wang, Heather E. Durfee, Yasmeen Hassan, Stephanie M. Bumbaru, Anthony Barbu, Alycen Wiacek, Dao-Qi Zhang

## Abstract

Amacrine cells (ACs) are retinal interneurons that regulate synaptic transmission from bipolar cells to retinal ganglion cells (RGCs) and play essential roles in object motion detection, contrast sensitivity, and light adaptation. A subtype of GABAergic ACs identified using a tyrosine hydroxylase (TH) promoter–driven green fluorescent protein (GFP) mouse line has been termed TH2 amacrine cells (TH2-ACs). Although TH2-ACs contribute to the feature selectivity of object-motion signals in the adult retina, their functional properties during early postnatal development remain unclear. Using genetic mouse models, electrophysiology, immunohistochemistry, and calcium imaging, we show that TH2-ACs exhibit spontaneous rhythmic depolarizations during development. In the first postnatal week, these depolarizations were abolished by acetylcholine receptor antagonists, indicating that TH2-ACs are excited by starburst amacrine cells (SACs) via spontaneous cholinergic retinal waves. During the second postnatal week, rhythmic depolarizations persisted but were blocked by glutamate receptor antagonists, demonstrating that TH2-ACs are subsequently driven by bipolar cells through glutamatergic waves. Calcium imaging further revealed that this activity propagates across the TH2-AC network in a wave-like manner, potentially resulting in spatially and temporally patterned GABA release. Pharmacological blockades of GABA_A_ receptors significantly enhanced glutamatergic wave activity in SACs and RGCs, indicating that GABAergic signaling from TH2-ACs participates in exerting inhibitory control over retinal waves. Together, these findings identify TH2-ACs as active participants in the development of retinal wave circuits and suggest that this participation via GABA signaling could contribute to activity-dependent refinement of retinal circuits underlying object motion processing.

**Key points:**

- TH2 amacrine cells are excited by starburst amacrine cells through cholinergic retinal wave activity during the first postnatal week.
- During the second postnatal week, TH2 amacrine cells are driven by bipolar cells via glutamatergic retinal wave activity.
- The dense dendritic arborization of TH2 amacrine cells enables their participation in the propagation of both cholinergic and glutamatergic waves.
- Wave-like GABA release from TH2 amacrine cells contributes to the modulation of retinal wave activity through activation of GABA_A_ receptors.

## INTRODUCTION

Amacrine cells (ACs) are retinal interneurons that mediate signal transmission from bipolar cells to retinal ganglion cells (RGCs), contributing to visual functions such as light adaptation, contrast sensitivity, and object motion detection. One distinct subtype, visualized using tyrosine hydroxylase (TH) promoter-driven fluorescent reporter mouse lines, has been identified as type 2 catecholaminergic ACs or TH2-ACs (Brüggen et al., 2015; Zhang et al., 2004). Similar to type 2 catecholaminergic ACs in other species (Mariani and Hokoc, 1988; Tauchi et al., 1990; Törk and Stone, 1979), TH2-ACs are found in both the inner nuclear layer (INL), referred to as conventional TH2-ACs (cTH2-ACs), and the ganglion cell layer (GCL), referred to as displaced TH2-ACs (dTH2-ACs). Both cTH2-ACs and dTH2-ACs extend a single primary dendrite that arborizes within the middle of the inner plexiform layer (IPL), thereby positioning these cells to receive convergent input from ON and OFF bipolar cells (Brüggen et al., 2015; Knop et al., 2011; Zhang et al., 2004). Excitation from ON and OFF bipolar cells drives TH2-ACs, which in turn provide GABAergic inhibition to signal transmission from vesicular glutamate transporter 3-expressing amacrine cells (VG3-ACs) to object motion-sensitive retinal ganglion cells (W3-RGCs) (Brüggen et al., 2015; Kim and Kerschensteiner, 2017). Through this inhibitory pathway, TH2-ACs shape the feature selectivity of object motion signals transmitted to the brain (Kim and Kerschensteiner, 2017). Despite their well-defined role in mature retinal circuits, the developmental properties of TH2-ACs and their contributions to early retinal network activity remain unknown.

During early postnatal development, spontaneously coordinated activity known as retinal waves propagates across the retina (Blankenship and Feller, 2010; Kerschensteiner, 2016). In the mammalian retina, these waves are classified into three distinct stages. Stage I occurs before birth and is mediated by electrical coupling through gap junctions, together with cholinergic signaling from mirror-symmetric ON and OFF starburst amacrine cells (SACs) (Voufo et al., 2023). After birth, Stage II cholinergic retinal waves predominate during the first postnatal week. In the second postnatal week, cholinergic waves are subsequently replaced by Stage III glutamatergic waves initiated by bipolar cells (Akrouh and Kerschensteiner, 2013a; Arroyo and Feller, 2016; Xu et al., 2016; Zhang et al., 2016; Zhou, 1998). Glutamatergic retinal waves subside or disappear after eye opening, coinciding with the onset of visually evoked activity (Wong et al., 1993). Although the cellular sources of cholinergic and glutamatergic waves are relatively well understood, it remains unclear how these waves drive the activity of other AC types and whether those ACs contribute to the propagation of cholinergic or glutamatergic waves, and vice versa.

TH2-ACs are an ideal cell type for addressing these research questions, owing to their distinctive morphology and functional properties. Their dendritic stratification in the middle of the IPL positions them between the dendritic bands of ON and OFF SACs, suggesting that TH2-ACs are anatomically suited to interact with cholinergic circuits during development (Brüggen et al., 2015; Mariani and Hokoc, 1988; Zhang et al., 2004). However, it remains unclear whether TH2-ACs express acetylcholine (ACh) receptors that enable them to respond during Stage II cholinergic waves. In addition, TH2-ACs receive inputs from both ON and OFF bipolar cells in the mature retina (Huang et al., 2022; Kim and Kerschensteiner, 2017; Knop et al., 2011), implying that they may also be activated during glutamatergic waves in the second postnatal week. This hypothesis, however, has not yet been tested.

TH2-ACs are a subset of GABAergic interneurons (Knop et al., 2011). Their release of GABA release could be driven by graded potentials and action potentials. Previous work has demonstrated that TH2-ACs are non-spiking neurons in adults (Knop et al., 2011). However, it is unclear whether TH2-ACs are incapable of generating action potentials at all, or whether they can generate action potentials, similar to SACs, during development but lose this ability in adulthood (Yan et al., 2020; Zhou and Fain, 1996). Additionally, GABA has an excitatory effect during early development due to the high intracellular chloride concentration in immature neurons; however, this effect becomes inhibitory in late development and adulthood (Leitch et al., 2005; Wang et al., 2007; Zhang et al., 2006). Despite this difference, endogenous GABA appears to suppress cholinergic and glutamatergic wave activity during early and late development, respectively (Hennig et al., 2011a; Wang et al., 2007; Zhou and Xu, 2015). To understand the physiological function of TH2 ACs, it is worth revisiting whether GABA shapes the strength and spatial patterns of glutamatergic waves immediately after the transition from its excitatory to inhibitory roles.

In this study, we combined genetic mouse models with electrophysiological recordings, immunohistochemistry, and calcium imaging to address several key questions: Do SACs excite TH2-ACs during cholinergic waves? Are TH2-ACs driven by bipolar cells during glutamatergic waves? Can TH2-ACs generate action potentials to trigger GABA release? Is the TH2-AC dendritic network sufficient to support or propagate retinal waves? Does GABA released from TH2-ACs modulate retinal wave propagation? We found that TH2-ACs exhibit spontaneous, rhythmic cholinergic and glutamatergic depolarizations, occasionally accompanied by action potentials, and that these depolarization-driven calcium signals can propagate across the TH2-AC network. These findings suggest that TH2-ACs may release GABA in a wave-like manner, providing feedback that modulates retinal wave propagation both locally and across the visual system.

## MATERIALS AND METHODS

### Mice

Four mouse lines were used in this study: TH promoter–driven green fluorescent protein (GFP) mice (stock No. RBRC02095, Riken, Japan) (Matsushita et al., 2002), TH-Cre;Ai162 calcium reporter mice, choline acetyltransferase (ChAT)-Cre;Ai95 calcium reporter mice, and wild-type mice (stock No. 000664, Jackson Laboratory, Bar Harbor, ME). TH-Cre;Ai162 mice were generated by crossing TH-Cre (stock No. 008601, Jackson Laboratory, Bar Harbor, ME) with Ai162 mice (stock No. 031562, Jackson Laboratory, Bar Harbor, ME). ChAT-Cre;Ai95 mice were produced by crossing ChAT-Cre (stock No. 006410, Jackson Laboratory, Bar Harbor, ME) with Ai95 mice (stock No. 028865, Jackson Laboratory, Bar Harbor, ME) (Yan et al., 2020). All strains were maintained on a C57BL/6 background. Animals were housed in a temperature- and humidity-controlled facility under a 12 h:12 h light–dark cycle with ad libitum access to food and water. Breeding pairs were established, and the date of birth was designated as postnatal day 0 (P0). Both male and female neonatal mice (P4-P12) were used. All animal procedures, including housing, breeding, and animal euthanasia for tissue collection, were performed in accordance with the National Institutes of Health Guide for the Care and Use of Laboratory Animals and were approved by the Institutional Animal Care and Use Committee (IACUC) of Oakland University.

### Electrophysiology

We performed patch-clamp electrophysiological recordings from individual TH2-ACs in flat-mounted retinas. This procedure involved retinal dissection, continuous perfusion with temperature control, identification and visualization of TH2-ACs, and subsequent cell patching, data acquisition, and analysis. For retinal dissection, pups were euthanized either by decapitation or by an overdose of CO₂ followed by cervical dislocation. Eyeballs were then enucleated and transferred to a Petri dish containing Ames’ medium bubbled with 95% O₂/5% CO₂. Retinas were dissected under dim red light and placed in a recording chamber with the ganglion cell layer facing upward. The chamber was mounted on the stage of an upright fluorescence microscope (Axio Examiner, Zeiss, Oberkochen, Germany). Oxygenated Ames’ medium was continuously superfused at 1–3 ml/min, and the temperature was maintained at 32–34 °C using a temperature controller (TC-344A, Warner Instruments, Hamden, CT).

TH2-ACs were identified in *TH*::GFP retinas using a fluorescein isothiocyanate (FITC) filter set. GFP images were acquired with a digital camera (AxioCam MR3, Zeiss) and simultaneously displayed on a computer monitor via AxioVision software. After locating and marking GFP-labeled TH2-ACs, we switched from the fluorescence filter set to infrared differential interference contrast (IR-DIC) optics and monitored the targeted TH2-ACs under transmitted light for patch-clamp recording. In addition, RGCs were visualized directly under IR-DIC optics in wild-type retinas for patch-clamp recording.

Prior to patch-clamp recording, we first approached the retina with an empty glass pipette under IR-DIC optics. This pipette was used to gently break the inner limiting membrane (ILM) over the ganglion cell layer. After creating an opening in the ILM, the empty pipette was replaced with a new recording pipette filled with intracellular solution containing (in mM): 120 K-gluconate, 5 NaCl, 4 KCl, 10 HEPES, 2 EGTA, 4 Na₂-ATP, and 0.3 Mg-GTP. With this solution, pipette resistance was approximately 10 MΩ. The recording pipette was advanced through the opening in the ILM while applying gentle positive pressure. Once the pipette tip approached the target cell membrane, positive pressure was released, and slight negative pressure was applied to form a gigaohm seal. Additional negative pressure was then applied to rupture the membrane and establish the whole-cell configuration in voltage-clamp mode. After achieving whole-cell access, we switched to current-clamp mode to record the resting membrane potential and spontaneous activity in gap-free mode using an Axopatch 700B amplifier (Molecular Devices, San Jose, CA). To assess membrane properties, a series of current steps (−100 pA to +100 pA, 20 pA increments, 500 ms duration) was injected into the cell. Data were acquired using a Digidata 1440A digitizer and Clampex 10.7 software (Molecular Devices).

Data were analyzed offline using Clampfit 10.7 (Molecular Devices, Sunnyvale, CA). Spontaneous depolarization events in TH2-ACs and RGCs were counted manually over the total recording duration, and their frequency (events/min) was calculated by dividing the number of events by the recording time. Action potentials of RGCs were filtered using a 5 Hz low-pass filter to reveal spontaneous depolarizing graded potentials. The rising time and total charge for each RGC graded potential were measured using the Statistics function of Clampfit.

### Calcium imaging and imaging analysis

The procedures for animal euthanasia, retinal dissection, perfusion, recording medium, and fluorescent imaging were the same as those used for electrophysiological recordings. Images were acquired using a 20X water-immersion objective and a digital camera (XM10, Olympus, Tokyo, Japan) mounted on an upright fluorescence microscope (BX51WI, Olympus, Tokyo, Japan). Using CellSens software, time-lapse images were collected from retinas of TH-Cre;Ai162 mice. Ca²⁺ transients were quantified offline in individual TH2-ACs using CellSens software, and transient traces were constructed by plotting integrated fluorescence intensity as a function of time.

Using the same experimental procedures and Ca²⁺ imaging methods, time-lapse images were also acquired from SACs in retinas of ChAT-Cre;Ai95 mice. Ca^2+^ imaging videos were converted to .avi (Audio Video Interleave) files, which were then processed in MATLAB to generate heatmap videos and enable automated quantitative analysis. Video frames were normalized by the median of the video sequence to remove static structural differences within each video. Changes in illumination within each video were corrected using a frame-wise illumination estimate, which was smoothed over time using a robust weighted linear least squares filter and applied as a multiplicative normalization factor. Following this background and illumination correction, pixel intensities were converted to Z-scores with Z-scores greater than 2.8 classified as wave activity, resulting in a binarized video of wave activity. A region propagation algorithm was then used to detect each wave and track it across the field of view (FOV), generating heatmap videos to visualize wave propagation. By tracking the centroid and size of the region through each frame, wave quantity, direction, area, frequency, and speed were saved for each video. The wave area as a percentage of the FOV was also computed, allowing normalization across waves that exceeded the imaging field. Wave frequency was measured as the total number of waves per video divided by the video duration in minutes.

The same set of converted .avi files was also analyzed using ImageJ to quantify wave-associated fluorescence intensity. For each FOV containing a wave, a region of interest (ROI) within the wave was selected to measure peak fluorescence intensity (F1). An identically sized ROI was then placed at the same location in a preceding FOV without a wave to measure background fluorescence intensity (F_0_). Relative fluorescence intensity was calculated as (F_1_ − F_0_) / F_0_.

### Immunohistochemistry

Immunohistochemistry was performed to label pan-NaV sodium channels on TH2-ACs using the protocol published by Mojumder et al., with minor modifications (Mojumder et al., 2007). Eyeballs from P10 *TH*::GFP mouse pups were harvested, fixed in 4% paraformaldehyde (PFA) for 2 h, embedded in OCT, and cryosectioned into 14-µm vertical sections mounted on microscope slides. Sections were rinsed in 1× phosphate-buffered saline (PBS) and subjected to a two-step antigen retrieval protocol. The protocol included incubation in Tris–EDTA–Tween antigen-retrieval buffer at 70 °C for 15 min to reverse PFA-induced protein cross-linking and increase the availability of pan-NaV epitopes for antibody binding, followed by incubation in the same buffer at room temperature for 35 min to allow protein renaturation and prevent tissue drying. After antigen retrieval, sections were rinsed in 1× PBS and 70% methanol (MeOH). To reduce autofluorescence induced by PFA fixation and heat-mediated antigen retrieval, sections were treated with 2% sodium borohydride (NaBH₄) for 2 min, followed by sequential rinses in 70% MeOH, deionized water, and PBS. Sections were then blocked in 5% bovine serum albumin (BSA) with 0.5% Triton X-100 in PBS for 30 min and incubated overnight at room temperature with primary antibodies containing rabbit anti–pan-NaV (1:500, Invitrogen, PA5-104465) and goat anti-GFP (1:1000, Invitrogen, PA5-143588). After rinsing with 1× PBS, sections were incubated for 1 h at room temperature with secondary antibodies, including Alexa Fluor 594 donkey anti-rabbit (1:500, Invitrogen, A21207) and Alexa Fluor 488 donkey anti-goat (1:500, Invitrogen, A32814). Finally, sections were rinsed and mounted using Vectashield Hardset Antifade Mounting Medium. Images were acquired on a Zeiss LSM 900 confocal microscope using a 63× objective and analyzed with ZEN Blue v3.5 software (Zeiss).

### Drugs

Hexamethonium bromide (HEX), NBQX (2,3-dioxo-6-nitro-1,2,3,4-tetrahydrobenzo[f]quinoxaline-7-sulfonamide), D-AP5 [D-(−)-2-amino-5-phosphonopentanoic acid], tetrodotoxin (TTX), and SR-95531 (Gabazine; 6-imino-3-(4-methoxyphenyl)-1(6H)-pyridazinebutanoic acid) were obtained from Hello Bio Inc. (Princeton, NJ, USA). Stock solutions were stored frozen and diluted into the extracellular medium immediately before use.

### Statistical analysis

Two independent groups with normally distributed data but unequal sample sizes were compared using an unpaired *t*-test with Welch’s correction. Paired *t*-tests were performed to compare the means of two related groups. When the data were not normally distributed, a nonparametric Mann–Whitney *U* test was used. Data are presented as the mean ± standard error of the mean (SEM). A *p* value < 0.05 was considered statistically significant.

## RESULTS

TH2-ACs are not immunoreactive to TH antibodies in the wild-type mouse retina; however, they can be reliably visualized using several transgenic reporter mouse lines (Brüggen et al., 2015; Huang et al., 2022; Newkirk et al., 2013; Vuong et al., 2015; Zhang et al., 2004). In this study, we used two such lines to target TH2-ACs during development. The first was the *TH*::GFP mouse line, in which TH2-ACs express GFP (Brüggen et al., 2015). This model enabled direct visualization of TH2-ACs for patch-clamp recordings, allowing us to assess their spontaneous electrical activity. The second model was generated by crossing TH-Cre mice with the Ai162 GCaMP6s reporter line. The resulting TH-Cre;Ai162 mice permitted real-time monitoring of spontaneous calcium transients in TH2-ACs during early postnatal development. In both models, a subset of dopaminergic amacrine cells (DACs) was also labeled with weak GFP intensity. TH2-ACs were readily distinguished from DACs by their high GFP intensity, smaller somata, single primary dendrites, and dendritic stratification in the middle sublamina of the IPL.

### TH2-ACs exhibit spontaneous rhythmic depolarization in the first postnatal week

The somata of TH2-ACs are located in the INL and are also displaced in the GCL (Fig. 1A). Despite their different somatic positions, both cTH2-ACs and dTH2-ACs arborize within the middle layers of the IPL. To determine their developmental properties, we began to perform whole-cell current-clamp recordings from dTH2-ACs in flat-mounted retinas isolated from *TH*::GFP mice at P5-6, as these cells are readily accessible in the ganglion cell layer (Fig. 1A). All 11 dTH2-ACs recorded exhibited spontaneous rhythmic slow depolarizations (Fig. 1B). The average frequency of these depolarizations was 0.45 ± 0.06 events/ min (n = 11). To assess the strength of these events, we measured their amplitude and duration. The mean peak amplitude was 11.19 ± 1.68 mV (n = 11), and the mean duration was 7.13 ± 0.26 s (n = 11).

**Figure 1.**
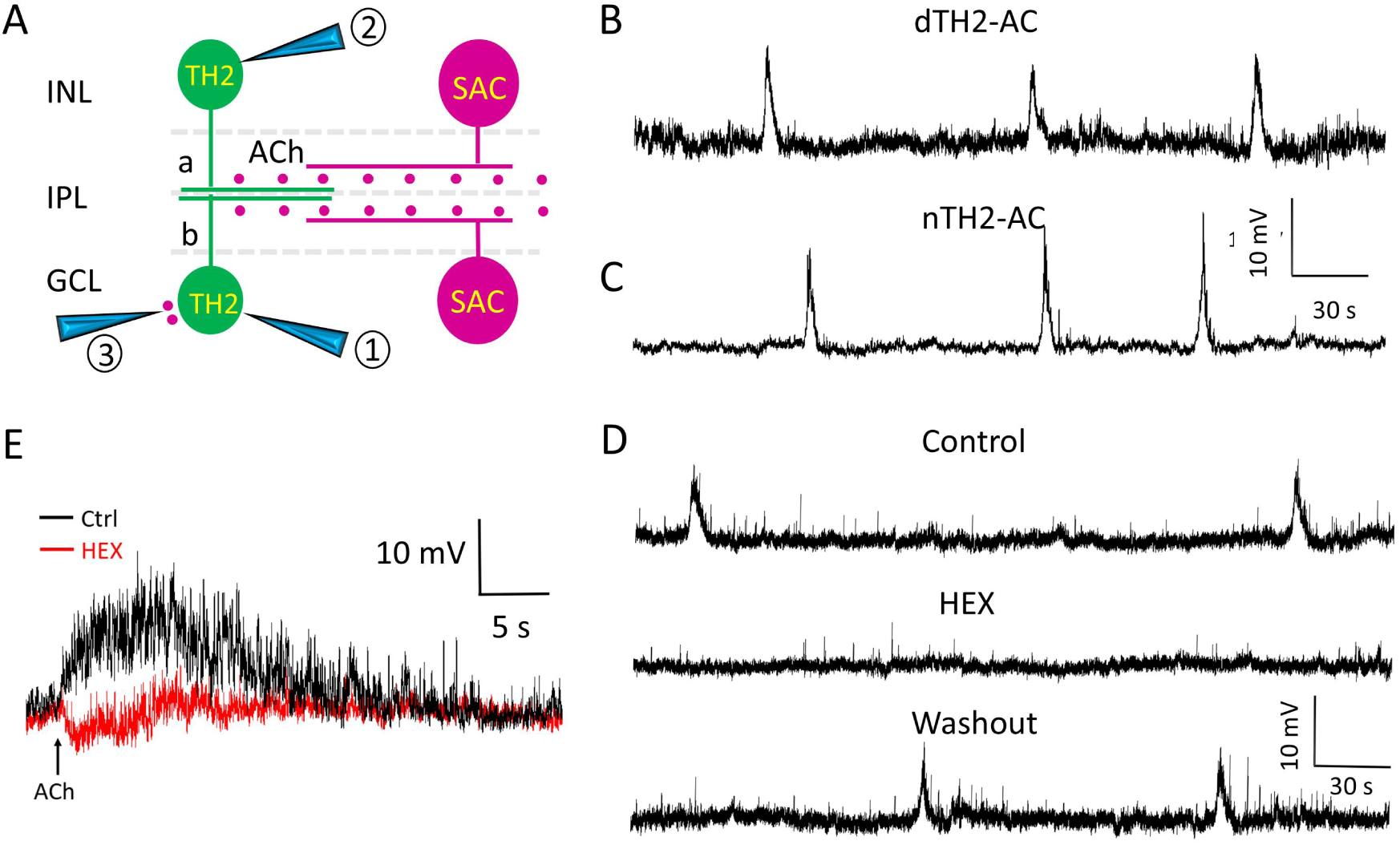
TH2-ACs are driven by SACs at the cholinergic wave stage. Whole-cell current-clamp recordings were obtained from cTH2-ACs and dTH2-ACs in flat-mount retinas isolated from *TH*::GFP transgenic mice at P5–P7. **A.** Schematic diagram illustrating the anatomical relationship between TH2-ACs and SACs. Positions 1 and 2 denote patch-clamp recording pipettes, and position 3 indicates the puff-application pipette. INL, inner nuclear layer; IPL, inner plexiform layer; GCL, ganglion cell layer; ACh, acetylcholine; a, sublamina a; b, sublamina b**. B.** Spontaneous rhythmic depolarizations recorded from a displaced TH2-AC (dTH2-AC). **C.** Spontaneous rhythmic depolarizations recorded from a cTH2-AC. **D.** Spontaneous rhythmic depolarizations in a dTH2-AC were abolished entirely by 100 µM hexamethonium (HEX), indicating dependence on cholinergic transmission. **E.** Local puff application of 1 mM ACh elicited a depolarization in a dTH2-AC, and pre-application of HEX suppressed this response, indicating functional expression of ACh receptors in TH2-ACs. Abbreviations defined in this figure are not be redefined if they appear in subsequent figures unless specified.

Because most TH2-ACs were located in the INL in both *TH*::GFP and TH-Cre;Ai162 mice, we next examined whether their physiological properties differed between cTH2-ACs and dTH2-ACs. Whole-cell recordings from cTH2-ACs in the INL were technically challenging due to their deeper location within the retina; nonetheless, we obtained recordings from four cells. All recorded cTH2-ACs exhibited spontaneous rhythmic depolarizations, similar to those observed in dTH2-ACs (Fig. 1C). The frequency of depolarizations (0.40 ± 0.04 events/min; n = 4) was not significantly different from that of dTH2-ACs (p = 0.727). Likewise, neither depolarization amplitude (8.53 ± 0.82 mV, n = 4) nor duration (6.76 ± 0.80 s, n = 4) differed significantly (p = 0.373 and p = 0.565, respectively). These results indicate that cTH2-ACs and dTH2-ACs exhibit comparable spontaneous activity patterns during early postnatal development.

#### Spontaneous depolarizations of TH2 ACs are mediated by cholinergic transmission

Both cTH2-ACs and dTH2-ACs ramify in the middle layers of the IPL, positioning them between the two cholinergic strata formed by the dendrites of ON and OFF SACs. This anatomical arrangement led us to hypothesize that TH2-ACs are activated by ACh released from SACs during the cholinergic retinal wave stage (Fig. 1A). To test this possibility, we bath-applied the ACh receptor antagonist hexamethonium (HEX, 100 μM) to the retina. Application of HEX (Fig. 1D, middle trace) abolished the spontaneous depolarizations observed in dTH2-ACs. This effect was reversible upon washout (Fig. 1D, bottom trace). Similar results were obtained in four additional cells. These findings suggest that spontaneous depolarizations in TH2-ACs originate from SACs.

Next, we tested whether TH2-ACs directly respond to ACh released from SACs during cholinergic retinal waves. We performed whole-cell recordings from dTH2-ACs and locally applied ACh using a glass pipette containing 1 M ACh positioned adjacent to the recorded cell (Fig. 1A). ACh was delivered between two spontaneous depolarization events to avoid overlap with ongoing rhythmic activity. In 7 out of 7 dTH2-ACs, local ACh application elicited depolarization (Fig. 1E). This ACh-evoked depolarization was abolished by HEX (Fig. 1E; n = 5). These results indicate that TH2-ACs express functional nicotinic ACh receptors and can directly respond to SAC-released ACh during cholinergic retinal waves.

### TH2-ACs exhibit cholinergic Ca^2+^ transients in a wave-like fashion across the retina

Our electrophysiological recordings demonstrate that individual TH2-ACs exhibit spontaneous depolarizations mediated by ACh. The next question is whether this depolarizing activity is confined to individual cells or propagates across neighboring TH2-ACs as a coordinated network event. To address this, we generated TH-Cre;Ai162 mice in which TH2-ACs express GCaMP6s, a high-sensitivity calcium indicator. In baseline images, TH2-ACs exhibited weak GFP fluorescence (Figure 2A, left). Within 2.58 seconds, calcium signals emerged in a cluster of TH2 cells in the lower-middle region of the field (Figure 2A, middle) and subsequently propagated toward the upper-left region (Figure 2A, right). The elevated fluorescence ultimately moved beyond the imaging frame. This spatiotemporal pattern is shown in full in Supplementary Video 1. Together, these imaging data demonstrate that calcium activity propagates across neighboring TH2-ACs in a wave-like manner, supporting the presence of a functionally interconnected TH2-AC network.

**Figure 2.**
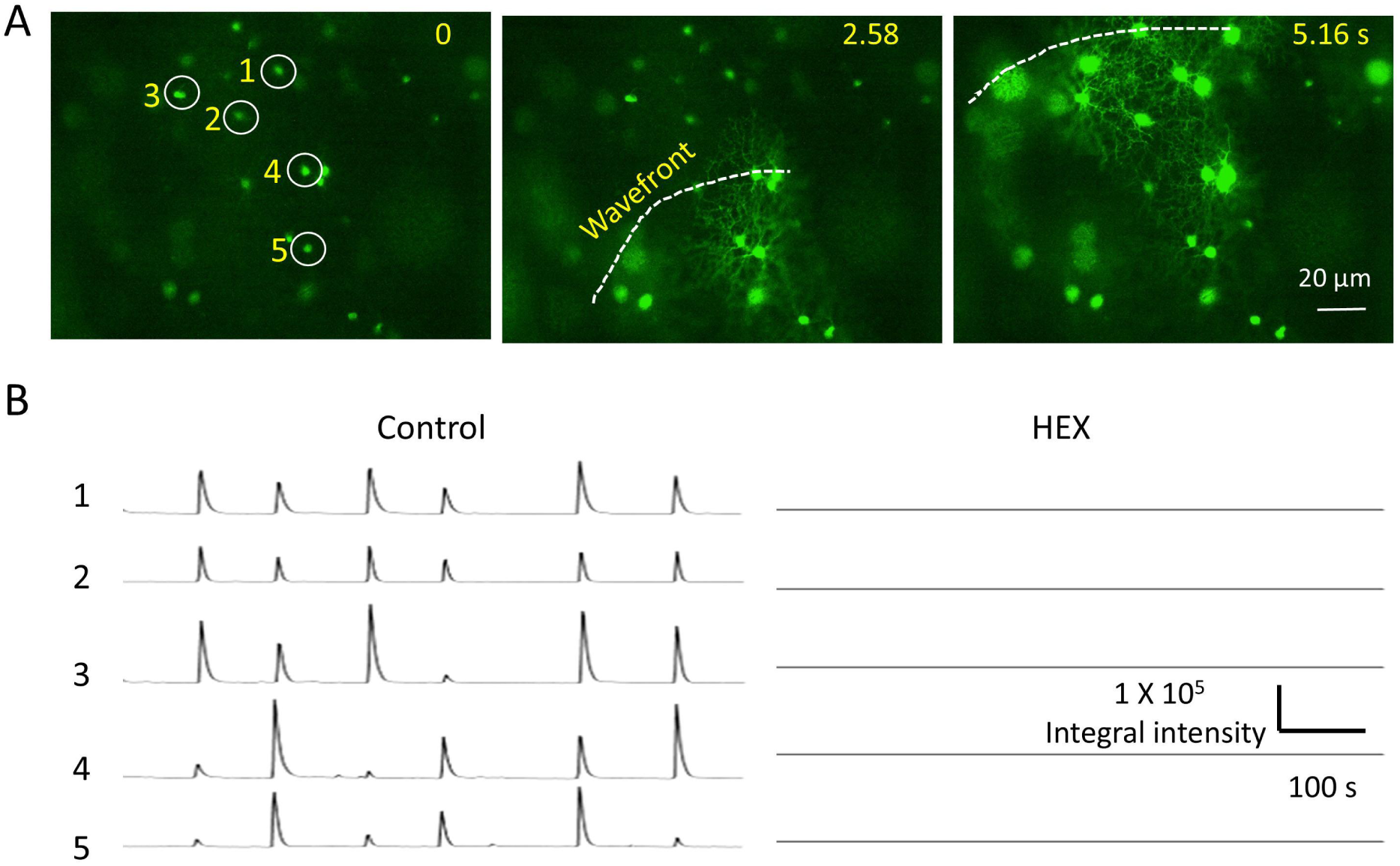
TH2-ACs participate in the propagation of cholinergic retinal waves. Time-lapse Ca²⁺ imaging was performed in flat-mount retinas isolated from TH-Cre;Ai162 mice at P6. **A.** The left image illustrates baseline Ca²⁺ signals in TH2-ACs. The middle and right images show a Ca²⁺ transient wave propagating from the bottom to the top of the imaging field. A video of this wave is provided in the supplementary materials. **B.** Five TH2-ACs were tracked, as indicated in the left image in panel A. Ca²⁺ transient traces were generated by plotting integrated fluorescence intensity as a function of recording time (10 min). The traces reveal spontaneous rhythmic activity (left panel). Application of HEX completely abolished this rhythmic activity (right panel), indicating that the Ca²⁺ transient waves are mediated by cholinergic transmission.

To determine whether these waves occurred periodically, we measured GFP intensity over time in five individual cells (Fig. 2A, left). Each cell displayed spontaneous, rhythmic Ca^2+^ transients, with a mean frequency of 0.57 events/min (Fig. 2B, left panel). To confirm that SACs are the source of these waves, we applied HEX, and it completely abolished wave activity (Fig. 2B, right panel). These results indicate that the dendritic networks of TH2-ACs are sufficient to support intercellular cholinergic wave propagation. Given that TH2-ACs are GABAergic, these findings suggest that GABA may also be released from these cells in a wave-like manner.

#### TH2-ACs exhibit spontaneous rhythmic depolarization in the second postnatal week

In the second postnatal week, cholinergic waves are gradually replaced by glutamatergic waves initiated by bipolar cells (Kerschensteiner, 2016). Given that TH2-ACs receive input from both ON and OFF bipolar cells (Fig. 3A), we hypothesized that they are excited by volumetric release of glutamate from bipolar cells during glutamatergic waves (Blankenship et al., 2009). To test this possibility, we used *TH*::GFP mice at P9 and P10. During those ages, cholinergic waves are almost entirely replaced by glutamatergic waves. Whole-cell current-clamp recordings were obtained from cTH2-ACs and dTH2-ACs in flat-mount retinas (Fig. 3A). We found that both subtypes exhibited spontaneous rhythmic depolarizations (Fig. 3B,C). Unlike ACh-mediated depolarizations, which exhibited a single peak, some depolarizations during this period showed more than one peak (Fig. 3B). This difference is discussed in the Discussion section.

**Figure 3.**
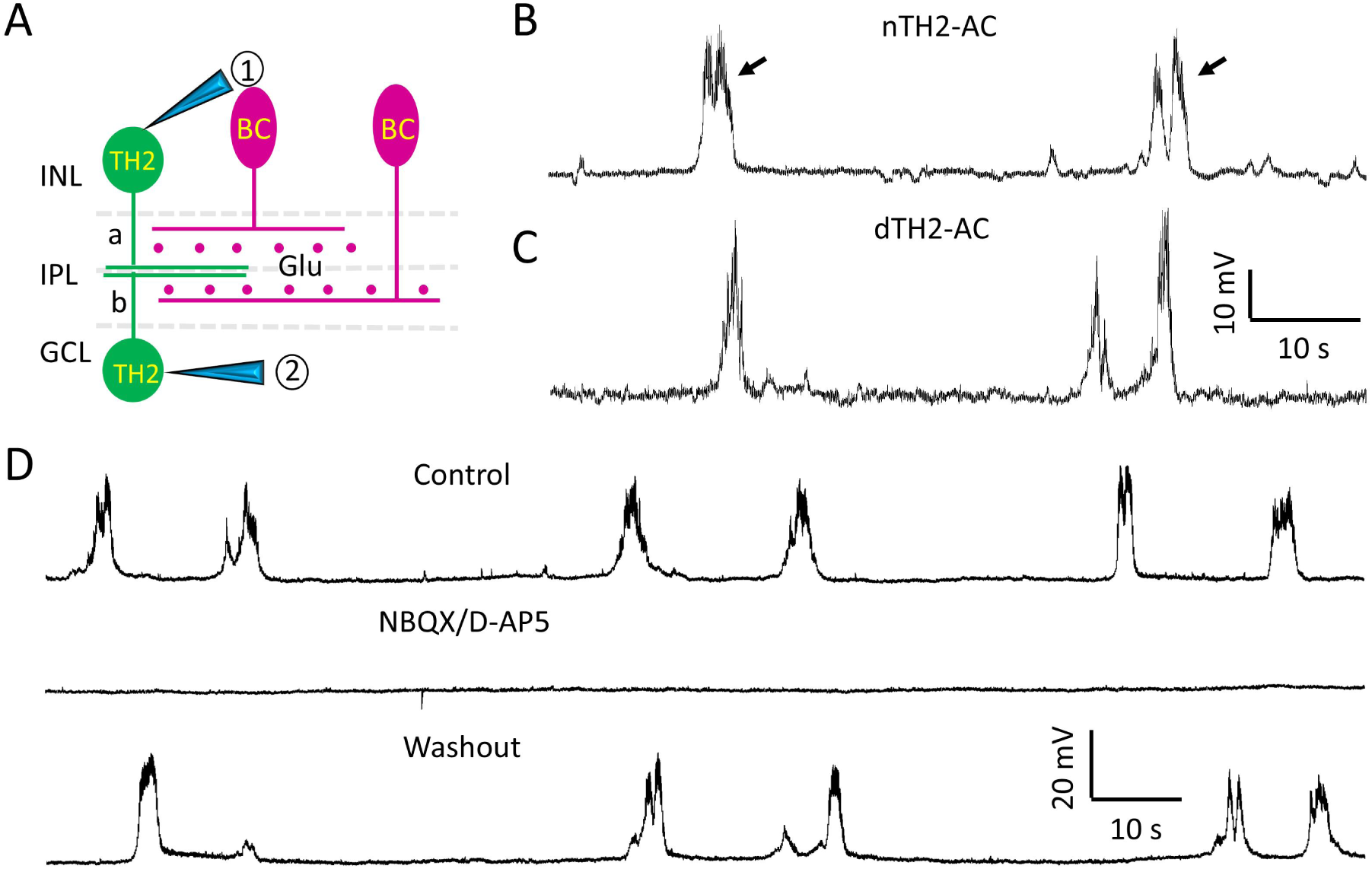
TH2-ACs are excited by bipolar cell input at the glutamatergic wave stage. Whole-cell current-clamp recordings were obtained from cTH2-ACs and dTH2-ACs in flat-mount retinas isolated from *TH*::GFP transgenic mice at P9–P10. **A.** Schematic diagram illustrating the anatomical relationship between TH2-ACs and bipolar cells (BCs). Positions 1 and 2 denote patch-clamp recording pipettes. Glu, glutamate. **B.** Spontaneous depolarizations recorded from a cTH2-AC. Arrows indicate the double-peaked depolarization waveform. **C.** Spontaneous depolarizations recorded from a dTH2-AC. **D.** Spontaneous rhythmic depolarizations in a dTH2-AC were abolished by a mixture of 50 µM NBQX and 50 µM D-AP4, indicating mediation of glutamatergic transmission.

Next, we performed further analysis to determine whether cTH2-ACs and dTH2-ACs exhibit similar or different levels of depolarization. We measured the frequency, amplitude, and duration of depolarization in cTH2-ACs and dTH2-ACs, respectively. The mean frequency did not differ significantly between dTH2-ACs (0.61 ± 0.09 events/min, n = 12) and cTH2-ACs (0.51 ± 0.11 events/min, n = 5; *p* = 0.539). The peak amplitude (24.66 ± 1.49 mV, n = 5) in cTH2-ACs was similar to that (21.85 ± 1.567 mV, n = 12) in dTH2-ACs (*p* = 0.304). Additionally, the mean duration in cTH2-ACs (4.66 ± 0.39 s, n = 5) was not significantly different from that in dTH2-ACs (6.11 ± 0.63 s, n=12; *p* = 0.173). Collectively, these findings indicate that cTH2-ACs and dTH2-ACs are similarly excited by bipolar cells during glutamatergic waves.

To confirm whether the spontaneous depolarizations of TH2-ACs are mediated by glutamate released during glutamatergic waves, we applied the glutamate receptor antagonists NBQX (50 μM) and D-AP5 (50 μM). Application of these antagonists completely abolished spontaneous depolarizations in dTH2-ACs (Fig. 3D, *n* = 4). This blockade was reversible upon washout, indicating that glutamatergic input from bipolar cells mediates the depolarization.

### Glutamatergic Ca^2+^ transients spread in a wave-like fashion across TH2 ACs

We next determined the extent to which glutamate-induced depolarization enhances intracellular calcium levels in TH2-ACs and whether this enhancement propagates between them, forming a wave at P9–P10. We monitored Ca²⁺ transients in TH2-ACs by time-lapse imaging of GCaMP6f expression in the middle of the IPL, where their dendrites are ramified. We observed that calcium wave activity propagated across the dendrites of TH2-ACs. As shown in Fig. 4A, a wave traveled from right to left across the imaging field. This spatiotemporal pattern is shown in full in Supplementary Video 2.

**Figure 4.**
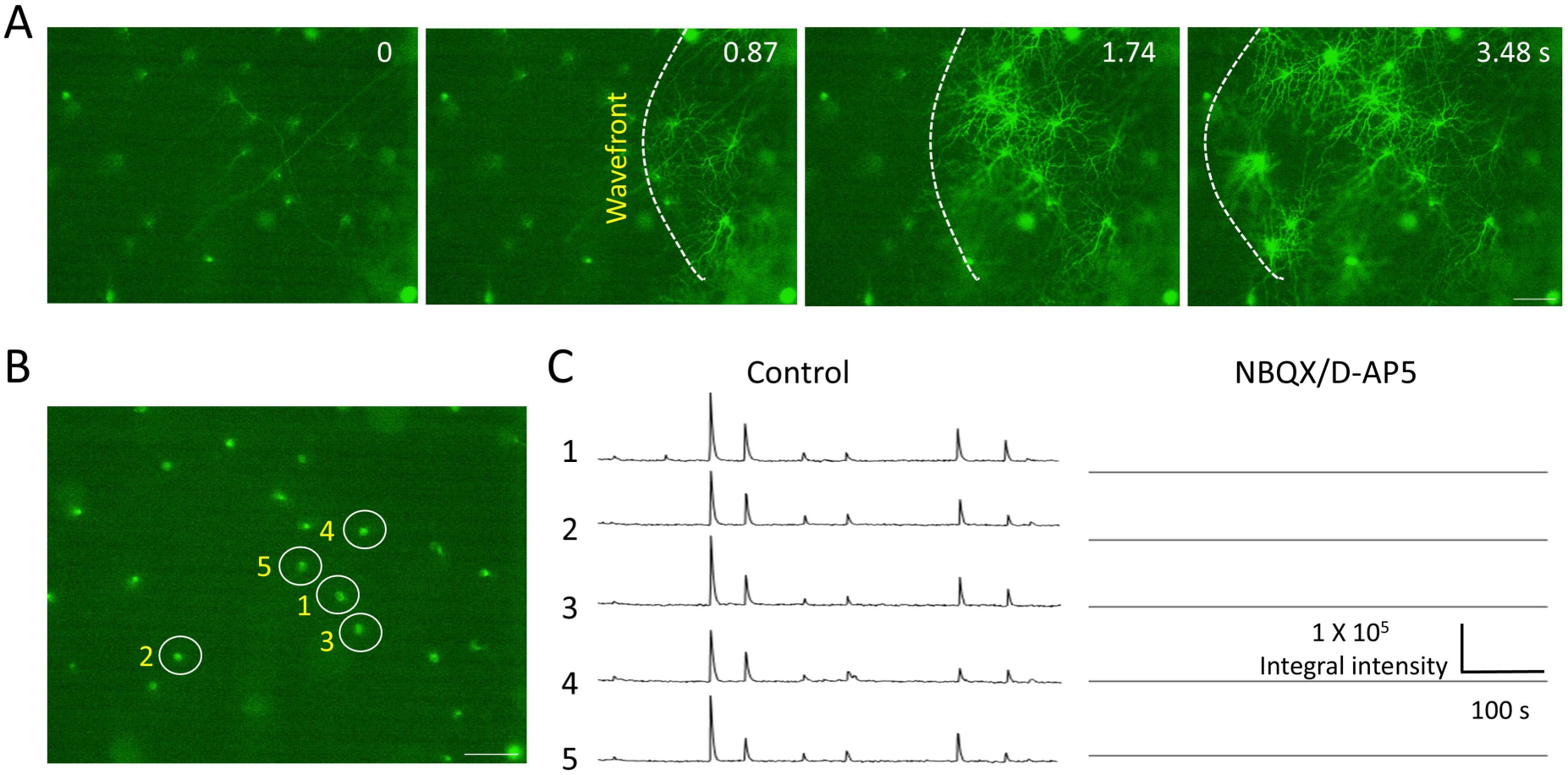
TH2-ACs participate in the propagation of glutamatergic retinal waves. Time-lapse Ca²⁺ imaging was performed in flat-mount retinas isolated from TH-Cre;Ai162 mice at P10. **A.** Imaging was focused within the IPL. The far-left image shows weak baseline Ca²⁺ signals in TH2-ACs. A Ca²⁺ transient wave first appeared on the right side of the imaging field (left middle image) and then propagated across the dendrites of TH2-ACs toward the left side (right middle and far-right images). A video of this wave is provided in the supplementary materials. **B.** Five TH2-ACs were tracked, as indicated by the numbered cells. **C.** Ca²⁺ transient traces were generated by plotting integrated fluorescence intensity as a function of recording time (10 min). The traces reveal spontaneous rhythmic activity (left panel). Application of NBQX/D-AP5 completely abolished this rhythmic activity (right panel), indicating that the Ca²⁺ transient waves are mediated by glutamatergic transmission. Scale bars in A and B: 50 µm.

To confirm that the calcium transient wave in TH2-ACs is driven by glutamatergic signaling, we traced five cells over time (Fig. 4B, left image), both before (Fig. 4C, middle trace panel) and during NBQX and D-AP5 application (Fig. 4C, right trace panel). We found that blocking AMPA and NMDA glutamate receptors eliminated Ca²⁺ transient propagation between TH2-ACs. The same results were observed in 7 additional mice. Taken together, these results suggest that TH2-ACs participate in glutamatergic retinal waves, which presumably drive TH2-ACs to release GABA in a wave-like manner.

#### Action potentials occasionally accompany spontaneous depolarizations of TH2-ACs

A previous study demonstrates that TH2 ACs do not generate action potentials in the adult retina (Knop et al., 2011). However, we observed a single action potential superimposed on spontaneous graded depolarizations in 8 of 30 TH2-ACs recorded between P5 and P11. An example is shown in Figure 5A, where an action potential is superimposed on a graded depolarization (see the insect in Fig. 5A). This finding suggests that TH2-ACs can generate action potentials, contrary to the previous report (Knop et al., 2011). To further explore this capability, we performed current injections on each recorded TH2-AC. We found that almost all cells exhibited current-induced action potentials, as shown in Figure 5B (left panel). These action potentials were usually single spikes rather than bursts, even when large positive currents (up to 100 pA) were applied to the cell. The application of the sodium channel blocker tetrodotoxin (TTX) completely blocked the action potentials, indicating that voltage-gated sodium channels mediate them. This pharmacological evidence was further validated by immunohistochemistry, demonstrating that TH2 amacrine cells express pan-NaV sodium channels at P10 *TH*::GFP mice (Fig. 5C, n = 4 mice). Overall, our results demonstrate that TH2-ACs can generate action potentials at least during development. Therefore, both graded depolarizations and action potentials can trigger GABA release in TH2-ACs, although action potentials are rarely initiated spontaneously with graded depolarizations.

**Figure 5.**
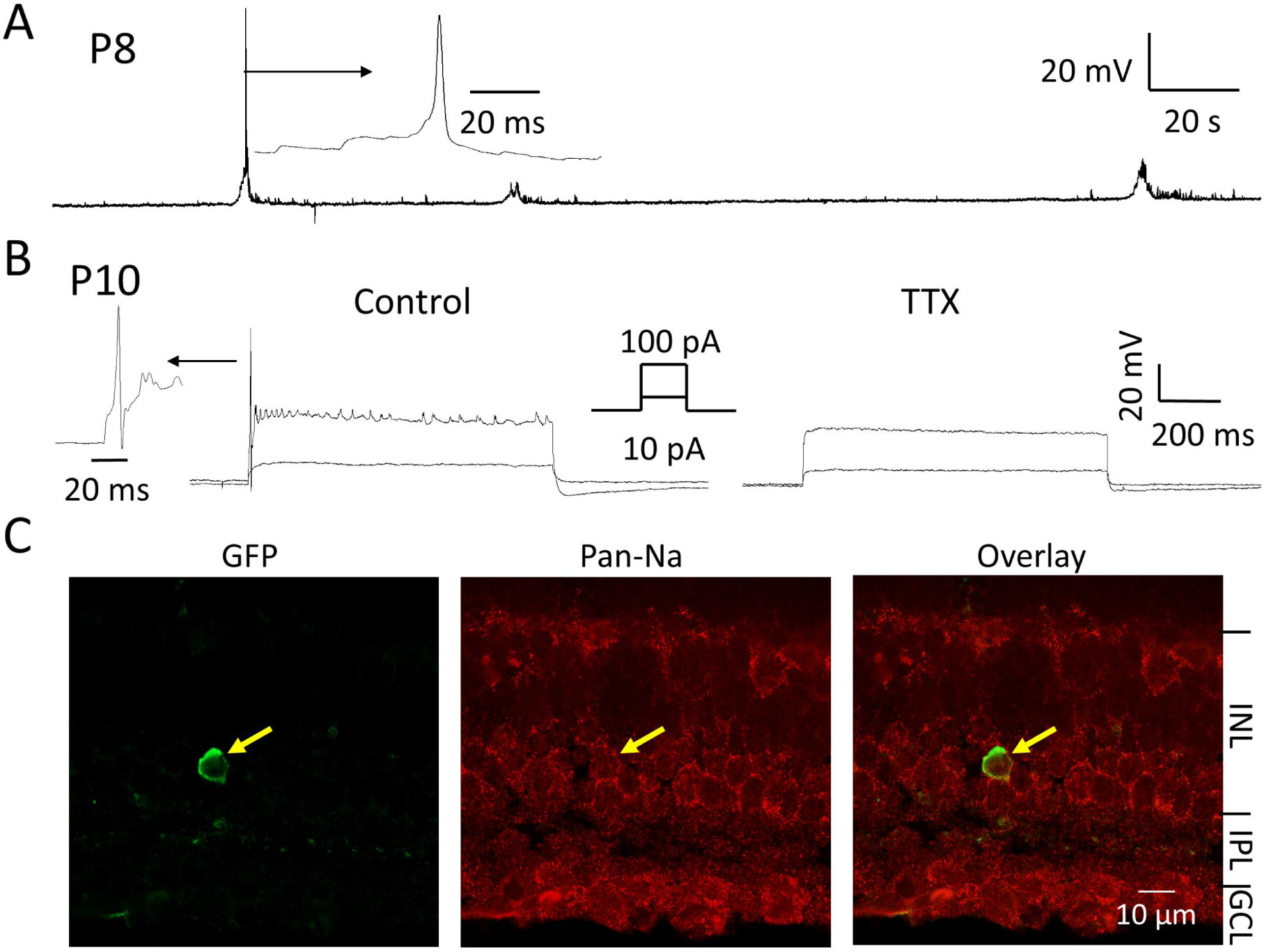
TH2-ACs generate sodium channel-mediated action potentials during development. Whole-cell current-clamp recordings were obtained from cTH2-ACs and dTH2-ACs in flat-mount retinas isolated from *TH::*GFP transgenic mice. **A.** A dTH2-AC at P8 exhibited spontaneous rhythmic depolarizations, with an action potential superimposed on one of the depolarizing events. The action potential waveform is shown at an expanded view in the inset. **B.** Step currents of 10 pA and 100 pA (shown in the inset) were applied to a dTH2-AC at P10. Injection of 100 pA induced a depolarization with a single action potential followed by a train of smaller spikes (left panel). Both the action potential and the spike train were completely abolished by the sodium channel blocker TTX (right panel). An expanded view of the action potential waveform is shown at the far left. **C**. The GFP signal in the vertical retinal section from P10 *TH*::GFP mice was enhanced by an anti-GFP antibody, revealing a cTH2-AC (left). The pan-NaV antibody labeled neurons across the retina (middle), including the cTH2-AC in this section (right).

#### Blockade of GABA_A_ receptors enhances glutamatergic waves monitored in SACs

Our electrophysiology and Ca^2+^ imaging data suggest that TH2-ACs participate in both cholinergic and glutamatergic retinal waves. We next determined whether they, along with other ACs, in turn modulate retinal wave activity through GABAergic signaling. GABA exhibits both excitatory and inhibitory functions during development, with a transition period occurring between P6 and P8 (Leitch et al., 2005; Wang et al., 2007; Zhang et al., 2006). To ensure that GABA functions in its inhibitory role, we used mice older than P9, a developmental stage at which glutamatergic waves predominate. Glutamatergic waves are generated in bipolar cells and propagate horizontally to nearby OFF SACs in the INL and vertically to ON SACs in the GCL (Fig. 6A). We have previously reported that ON and OFF SACs exhibited glutamatergic waves in ChAT-Cre;Ai95 mice (Yan et al., 2020). To determine the effect of endogenous GABA receptors on glutamatergic waves, we acquired time-lapse images of SACs in isolated retinas from ChAT-Cre;Ai95 mice before (Supplementary Video 3) and during application of 10 μM SR95531 (Supplementary Video 4). Calcium imaging videos were processed and visualized as heatmaps using a MATLAB script (see Methods for details). A representative heatmap in Fig. 6B (top) illustrates a retinal wave propagating from left to right over 3.87 s ( also see Supplementary Video 5). In the presence of SR95531, the spatial extent of the wave was markedly increased within the same time window, as shown in Fig. 6B (bottom) (also see Supplementary Video 6). Quantitative analysis revealed that wave area was approximately doubled during SR95531 treatment compared with control (Fig. 6C, *p* < 0.001). SR95531 also significantly increased the wave frequency (Fig. 6D, *p* < 0.05) and fluorescent intensity (Fig. 6E, *P* < 0.001). Notably, these increases persisted after a 20-min washout period, which we expected to be sufficient for complete drug washout (Fig. 6C–E). Together, these results indicate that GABA_A_ receptor blockade enhances the synchronization of glutamatergic retinal waves, resulting in increased wave area, frequency, and intensity.

**Figure 6.**
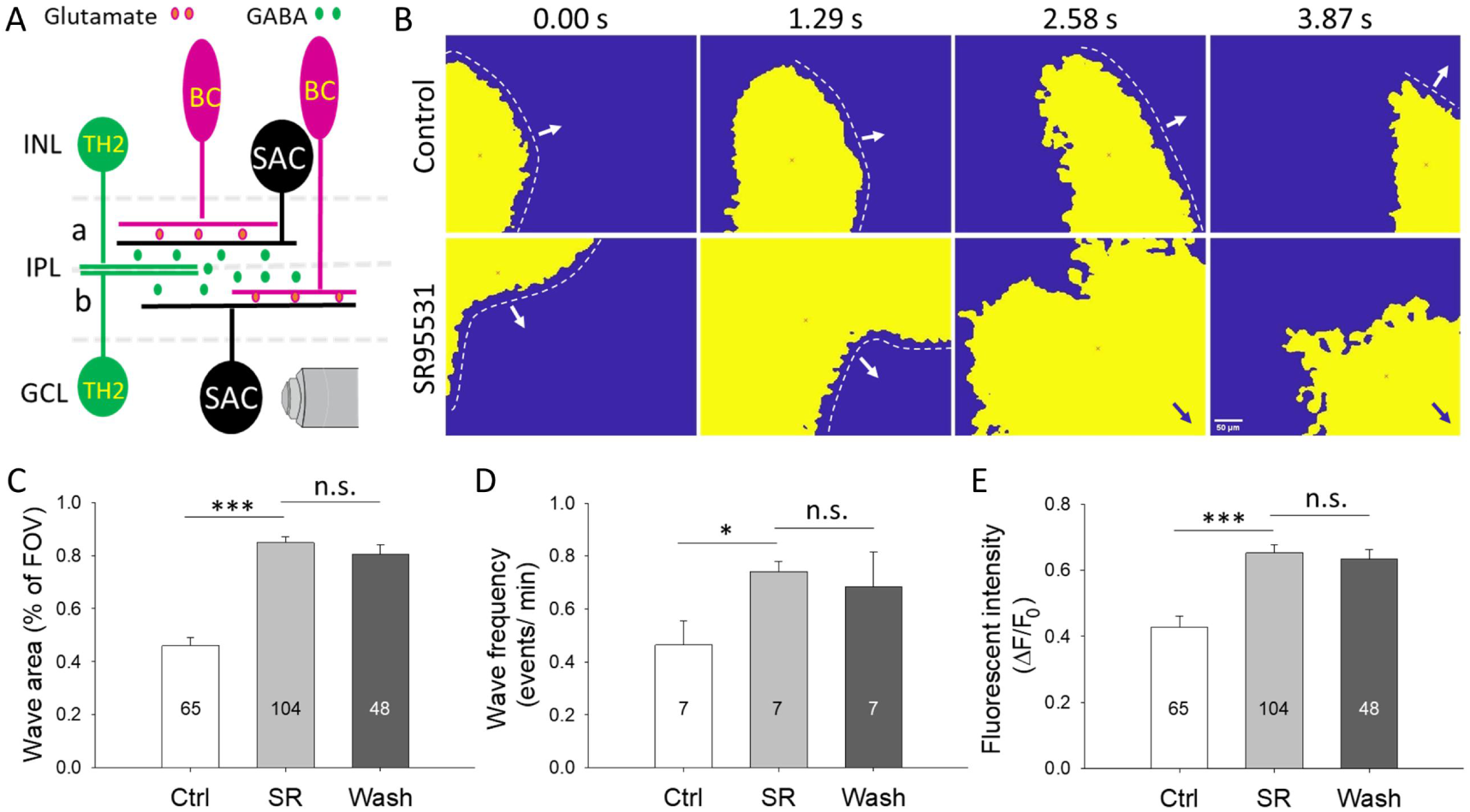
Blockade of GABA_A_ receptors enhances glutamatergic calcium waves in SACs. **(A)** Schematic diagram illustrating the anatomical relationships among TH2-ACs, bipolar cells (BCs), and SACs. Time-lapse calcium imaging was performed on SACs in ChAT-Cre;Ai95 mouse retinas at P9–P11. **(B)** Calcium imaging videos were processed using a custom MATLAB script to generate heat-map representations of wave activity. Representative video stills show glutamatergic waves under control conditions (top) and during application of the GABA_A_ receptor antagonist SR95531 (10 μM; bottom). **(C)** MATLAB-based analysis quantified wave area in control, during SR95531 application, and after washout (*n* = 65, 104, and 48 waves, respectively). Wave area was significantly increased during SR95531 treatment compared with control (*p* < 0.001), and this increase was not reversed following washout (*p* > 0.05). **(D)** The total number of waves recorded within a 10-min imaging period before, during, and after SR95531 application was calculated using MATLAB analysis. Wave frequency was significantly increased during GABA_A_ receptor blockade and did not recover after washout (n= 7 retinas). **(E)** Relative wave intensity was calculated as △F / F_0_ using ImageJ (see Methods) under control conditions (*n* = 65 waves), during SR95531 (*n* = 104 waves), and after washout (*n* = 48 waves). Blockade of GABA_A_ receptors non-reversibly enhanced glutamatergic wave intensity. Numbers within the bars in C, D, E indicate the sample sizes. * *p* < 0.05; \*\*\**p* < 0.001.

#### Blockade of GABA_A_ receptors facilitates glutamatergic wave-induced depolarizations in RGCs

Similar to SACs, RGCs are downstream neurons of bipolar cells, which are often used to study retinal waves (Fig. 7A). To assess the more dynamic effects of SR95531, we examined its impact on spontaneous rhythmic depolarization in RGCs during glutamatergic waves. Whole-cell current-clamp recordings of RGCs were performed between P9 and P12. As expected, RGCs exhibited spontaneous rhythmic depolarizations superimposed on bursts of action potentials. To isolate wave-induced grade potentials, action potentials were removed offline (See Methods). Compared to the control (Fig. 7B, top trace), SR95531 administration (10 µM) markedly increased the peak amplitude of depolarization (Fig. 7B, middle trace). On average, this increase was statistically significant (Fig. 7C) and consistent with the elevated wave intensity in SACs (Fig. 6E). In addition, the frequency of spontaneous depolarizations showed a modest increase, but this change did not reach statistical significance (Fig. 7D). This contrasts with the significant increase in wave frequency observed in SACs across the field of view, suggesting that waves recorded from individual RGCs do not capture all wave events occurring across the field of view as revealed by calcium imaging.

**Figure 7.**
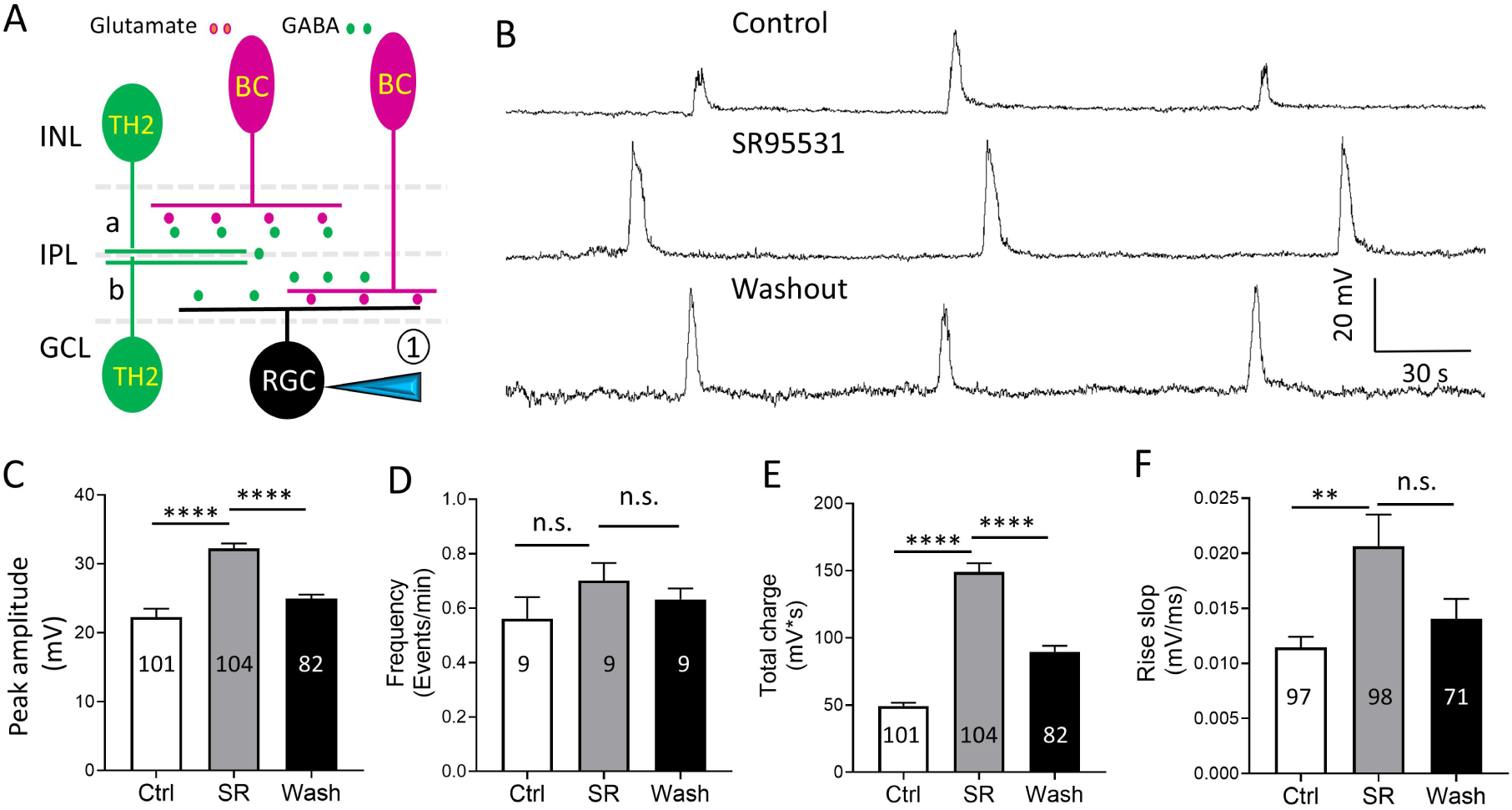
Blockade of GABA_A_ receptors enhances spontaneous rhythmic depolarization of RGCs at the glutamatergic wave stage. Whole-cell current-clamp recordings were obtained from RGCs in flat-mount retinas isolated from wild-type mice. **A.** Schematic diagram illustrating the anatomical relationship between TH2-ACs, bipolar cells (BCs), and RGCs. Position 1 denotes the patch-clamp recording pipette. **B.** RGCs exhibited a combination of spontaneous action potentials and rhythmic graded depolarizations. Action potentials were digitally filtered to reveal the underlying graded potentials (top panel). Application of 10 µM SR95531, a GABA receptor antagonist, increased the peak amplitude of the depolarization events (middle panel), and this enhancement was partially reversed 40 min after washout (bottom panel). **C.** Mean peak amplitude of depolarization events was significantly and reversibly enhanced by SR95531 (SR). **D.** The frequency of depolarization events tended to increase with SR95531, although the change did not reach statistical significance. **E.** SR95531 significantly increased the total charge of depolarization events, calculated as amplitude (mV) × depolarization duration (seconds). **F.** SR95531 significantly and reversibly increased the rising slope of depolarization events. Numbers within the bars indicate the number of events analyzed (C, E, F) or the number of RGCs analyzed (D). ** *p* < 0.01; \*\*\*\**p* < 0.0001.

To further quantify depolarization strength, we calculated the total charge per event by multiplying the depolarization amplitude by its duration. Total charge was approximately doubled in the presence of SR95531 (Fig. 7E), corroborating the increased wave area and intensity of glutamatergic waves observed in SACs (Fig. 6C,E). We next analyzed the slope of the depolarization rise phase before and during SR95531 application and found that SR95531 significantly increased the rising slope (Fig. 7F), indicating that GABA_A_ receptor blockade accelerates the temporal dynamics of RGC depolarization. A 20–40 min washout of the drug resulted in a significant reduction of the effect, but recovery remained incomplete (Fig. 7C, E, F). Together with the Ca²⁺ imaging data from SACs (Fig.6 C-E), these results suggest that blockade of GABA_A_ receptors could produce prolonged alterations in retinal wave activity. Together, our findings indicate that inhibition of GABA_A_ receptors markedly promotes both the strength and spatiotemporal extent of glutamatergic retinal waves, with effects that may persist due to activity-dependent plasticity within the developing glutamatergic wave network.

## Discussion

In this study, we found that cholinergic and glutamatergic waves drive GABAergic TH2-ACs during early development. These waves propagated within the TH2 AC network, suggesting that GABA release from them occurs in a wave-like manner. Pharmacological blockade of GABA_A_ receptors enhanced the strength and synchronization of glutamatergic waves, indicating that endogenous GABA shapes the spatiotemporal patterning of retinal waves. This regulation ultimately contributes to the refinement of neural circuits during visual system development.

### TH2-ACs and their cross-species type 2 catecholamine ACs

This study used two transgenic mouse models in which TH2-ACs were visualized throughout development, thereby enabling electrophysiological recordings and calcium imaging. These approaches allowed us to characterize the spatiotemporal dynamics of retinal wave activity in this subtype of ACs, which are essential for visual circuit development. Although our experiments were conducted in mice, the findings may offer broader insights into the developmental functions of type 2 catecholaminergic ACs in other mammalian species, such as rats, rabbits, and primates (Mariani and Hokoc, 1988; Tauchi et al., 1990; Törk and Stone, 1979). In these species, type 2 catecholaminergic ACs can be identified using TH immunolabeling, suggesting that they could co-release GABA and epinephrine. Type 2 catecholaminergic ACs exhibit somatic distributions and dendritic stratification patterns that closely resemble those of TH2-ACs. Given their structural and neurochemical similarities, TH2-ACs and type 2 catecholaminergic ACs across species may play similar roles in modulating early retinal network activity and refining visual circuits during development.

### Mutual interaction between TH2 ACs and SACs during development

TH2-ACs and SACs are two GABAergic amacrine cell subtypes with somata distributed in both the INL and GCL. Both cTH2-ACs and dTH2-ACs extend their dendrites into a single stratified band in the mid–inner plexiform layer. This band lies between the ON and OFF dendritic strata of SACs, positioning TH2-ACs to form reciprocal interactions with SACs. During cholinergic waves, SACs release ACh via volume transmission (Ford et al., 2012), and the diffusion of ACh to nearby TH2-ACs can excite these cells. Indeed, we observed robust wave-driven depolarizations and Ca²⁺ transients in TH2-ACs that were tightly correlated with cholinergic wave activity. Our physiological data indicate that TH2-ACs express functional ACh receptors and that ACh released from SACs directly activates these receptors to mediate the depolarization response. This is supported by similar cholinergic mechanisms observed in RGCs (Feller et al., 1996), suggesting that ACh receptors are broadly expressed in developing retinal neurons to ensure coordinated responses to cholinergic wave activity. Most importantly, both TH2-ACs and SACs release GABA, and reciprocal GABAergic inhibition between these cell types may strengthen the overall GABAergic regulation of retinal wave initiation, propagation, and termination.

### Inputs from bipolar cells to TH2-ACs during early development

In the mature retina, ON and OFF bipolar cells synapse onto TH2-ACs, resulting in both ON and OFF light responses in these cells, regardless of whether they are located in the INL or GCL (Huang et al., 2022; Kim and Kerschensteiner, 2017; Knop et al., 2011). During glutamatergic waves, it is postulated that ON bipolar cells initiate the wave, and the resulting glutamate spillover depolarizes neighboring GABAergic amacrine cells (Akrouh and Kerschensteiner, 2013b). These GABAergic ACs then provide inhibitory feedback onto OFF bipolar cells, hyperpolarizing them. When the glutamate spillover from ON bipolar cells subsides and the inhibitory input terminates, OFF bipolar cells undergo a rebound depolarization that triggers glutamate release from them. This sequence elicits the characteristic sequential ON-OFF responses in RGCs during Stage III glutamatergic waves (Akrouh and Kerschensteiner, 2013b; Kerschensteiner et al., 2009).

Our results reveal that glutamatergic waves also evoke robust depolarizations in TH2-ACs, with some cells exhibiting more than one depolarization peak. Such oscillatory depolarizations may reflect the circuit dynamics involving ON bipolar cells, intermediate GABAergic amacrine cells, and OFF bipolar cells. Notably, these intermediary GABAergic ACs could include TH2-ACs themselves. If so, ON bipolar cells would directly excite TH2-ACs, which in turn would provide GABAergic input to OFF bipolar cells, contributing to the delayed activation of OFF bipolar cells and generating the secondary peak of glutamatergic wave activity (Akrouh and Kerschensteiner, 2013b).

### Sufficient TH2-AC dendritic networks for wave propagation

The density of TH2-AC somata ranged from 33 to 205 cells/mm² across the retina, which is relatively low compared to that of SACs (∼400 cells/mm^2^) (Spix et al., 2016; Zhang et al., 2004). Despite their sparse somatic distribution, our Ca^2+^ imaging data show that the overlapping dendritic arbors from both cTH2-ACs and dTH2-ACs ensure nearly continuous coverage of the middle of IPL. This extensive dendritic field architecture enables TH2-ACs to form a functional network that supports the propagation of wave-driven activity. Thus, even with a lower somatic density, our data demonstrate that the dendritic organization of TH2-ACs provides a structural substrate sufficient for transmitting cholinergic and glutamatergic wave signals across them. In this way, TH2-ACs actively participate in the propagation of both cholinergic and glutamatergic waves. This enables TH2-ACs to provide continuous GABAergic feedback to nearby bipolar cells, SACs, and other amacrine cells. Together with other GABAergic ACs, this feedback dynamically regulates the amplitude, duration, and propagation speed of each wave event, ensuring proper wave initiation, propagation, and termination.

### Wave-induced graded membrane depolarizations and action potentials in TH2-ACs

Our results indicate that TH2-ACs begin to express functional ACh receptors and AMPA/NMDA-type glutamate receptors before eye-opening. In response to ACh or glutamate, activation of these receptors depolarizes cells in a form of graded potentials, as evidenced by our patch-clamp recordings, and increases intracellular Ca²⁺ levels, as demonstrated by calcium imaging. Elevated Ca²⁺ levels can trigger the release of neurotransmitters such as GABA and epinephrine onto nearby neurons, modulating their development. In addition, increased Ca²⁺ signaling may regulate their dendritic development, laminar organization, and synapse formation before eye opening. A future study will investigate the role of retinal waves in the structural development of TH2-ACs.

In addition, we occasionally observed action potentials superimposed on wave-induced depolarizations. This was surprising, as action potentials are not typically induced in adult TH2-ACs (Knop et al., 2011). We confirmed that most TH2-ACs exhibit current-injection-induced action potentials, suggesting that TH2-ACs can generate action potentials. This conclusion is further supported by immunohistochemical data showing that TH2-ACs express pan-sodium channels, and by the blockade of action potentials with TTX. We and others previously demonstrated that SACs only generate action potentials during development (Yan et al., 2020; Zhou and Fain, 1996). However, we did observe some TH2-ACs fire action potentials in adult retinas (data not shown). The overall data lead us to conclude that TH2-ACs can generate action potentials during development and throughout adulthood. This finding indicates that the action potentials superimposed on wave-induced depolarizations are physiologically meaningful and likely drive fast neurotransmitter release from TH2-ACs.

### The role of GABA_A_ receptors in the spatiotemporal dynamics of glutamatergic retinal waves

The larger glutamatergic waves observed in SACs under GABA_A_ receptor antagonism indicate that endogenous GABA desynchronizes the waves, as assessed by calcium imaging, and is consistent with previous multi-electrode array recordings from RGCs (Krishnan et al., 2025; Zhou and Xu, 2015). This desynchronization appears to alter the spatial pattern of the waves, which is essential for refining visual neural circuits. In addition, endogenous GABA likely attenuates wave strength, as evidenced by increased Ca²⁺ wave intensity in SACs and enhanced total charge of spontaneous depolarizations in RGCs when GABA_A_ receptors are blocked. This inhibitory feedback may negatively impact wave generation, propagation, and termination. Furthermore, endogenous GABA appears to slow wave propagation, as evidenced by an increased rising phase of spontaneous depolarization in RGCs and increased wave frequency in SACs upon GABA_A_ receptor blockade. This temporal dynamic effect underscores the role of GABAergic inhibition in regulating not only the spatial but also the temporal features of retinal wave activity, which are critical for proper developmental signaling in the visual system.

There are approximately 20 subtypes of GABAergic ACs, and TH2 ACs represent one of these subtypes. Our current study could not determine the specific contribution of TH2 ACs to the GABAergic regulation of retinal waves. However, due to their unique dendritic ramification in the middle layers of the IPL, they may play a more prominent role than other GABAergic ACs. GABAergic ACs can provide inhibitory feedback to bipolar cells, thereby limiting glutamate release during glutamatergic waves. In addition, they may act at GABA receptors on SACs or RGCs, thereby suppressing their responses to glutamate. These feedback, feedforward, and mutual interactions may integrate to shape the spatiotemporal properties of retinal waves. This integration may be subject to homeostatic plasticity (Hennig et al., 2011b), as blockade of GABA_A_ receptors produces prolonged alterations in retinal wave activity that persist beyond drug washout. Future studies will investigate the cellular and synaptic mechanisms underlying this persistence.

### Possible developmental role of the TH2-AC Network

We previously showed that TH2 ACs form a spatial gradient from the dorsal-nasal to the temporal-ventral retina, with lower density in the temporal-ventral region (Zhang et al., 2004). This distribution appears to parallel the temporal-to-nasal directional bias of retinal waves observed between postnatal days 8 and 10 (P8–P10) (Ge et al., 2021). We speculate that less TH2 AC density in the temporal retina lowers GABA levels, facilitating pacemaker excitability and wave propagation. In contrast, higher TH2 AC density in the nasal retina may enhance GABAergic inhibition, limiting excitability and wave propagation. This gradient may underlie the consistent temporal-to-nasal wave direction. Future studies are needed to test whether TH2 ACs contribute to this directional bias. If so, they may play a key role in shaping activity patterns essential for the development of visual circuits involved in optic flow processing.

In addition, TH2-ACs provide inhibitory input to the VG3 AC-W3 RGC circuit for object motion detection in the adult retina (Kim and Kerschensteiner, 2017). This inhibitory interaction suggests that, during development, GABAergic signaling from TH2 ACs may directly or indirectly influence the maturation of this circuit. Future studies will examine how TH2 ACs and retinal waves interact to shape the feature selectivity of object motion signals transmitted to the brain. Understanding this relationship may provide insight into how inhibitory interneurons contribute to the emergence of functional visual circuits during early postnatal development.

## Author contributions

Conceptualization: W.W. and D.-Q.Z.; data acquisition: W.W., H.D., S.B., A.B., and D.-Q.Z.; MATLAB script development: Y.H. and A.W.; data analysis: W.W., H.D., S.B., A.B., A.W., and D.-Q.Z.; manuscript drafting and revision: W.W., H.D., A.B., A.W., and D.-Q.Z. All authors have read and approved the final version of the manuscript, and all individuals who meet the criteria for authorship are listed.

---

**Supplementary Video 1.**
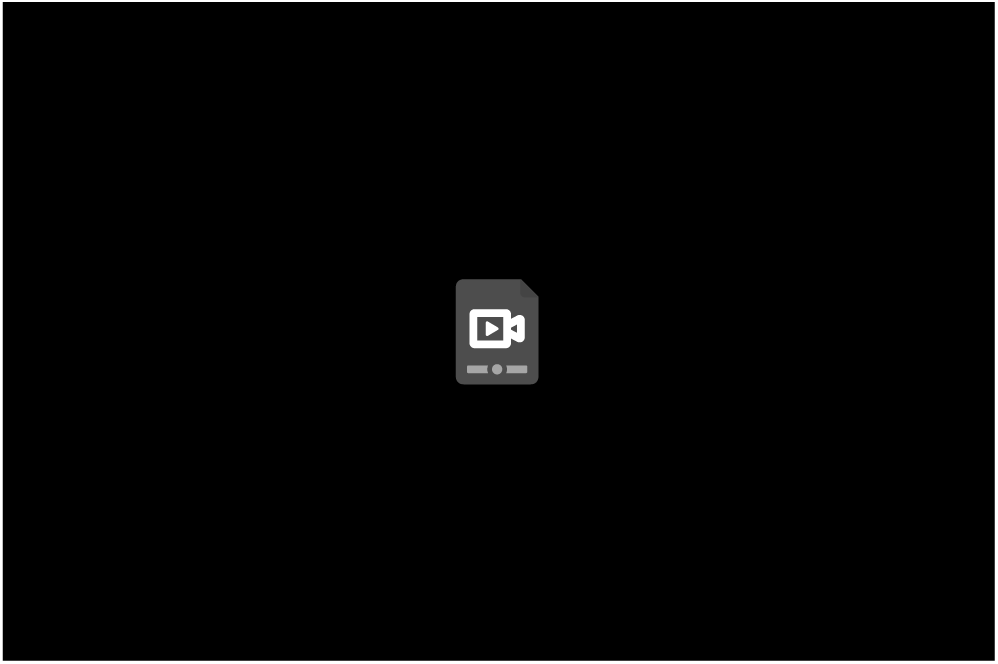
Corresponding time-lapse Ca^2+^ imaging video for Figure 2A.

**Supplementary Video 2.**
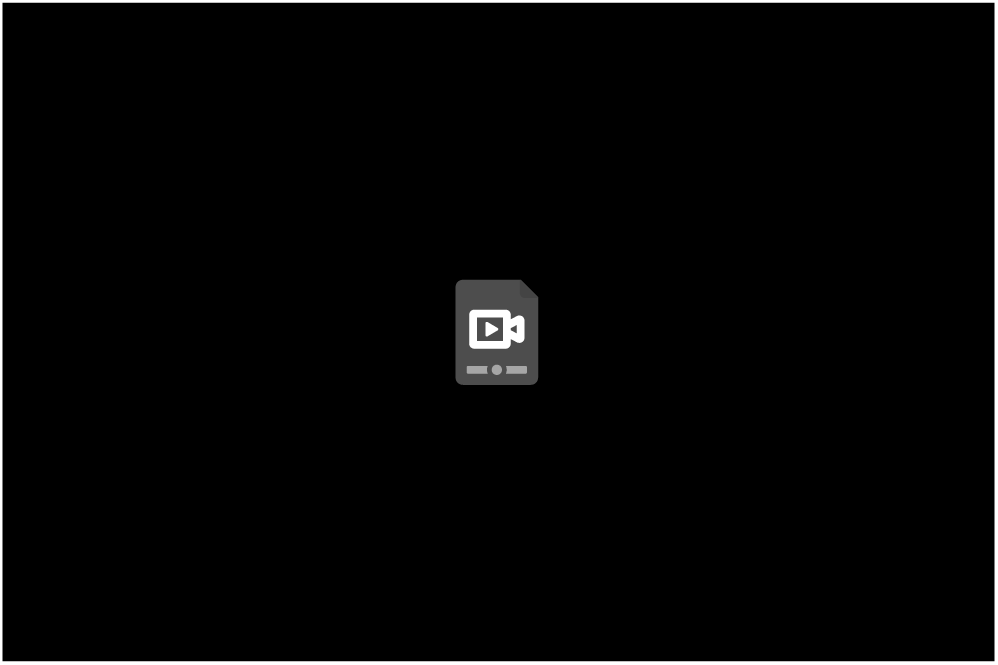
Corresponding time-lapse Ca^2+^ imaging video for Figure 4A.

**Supplementary Videos 3–4.**
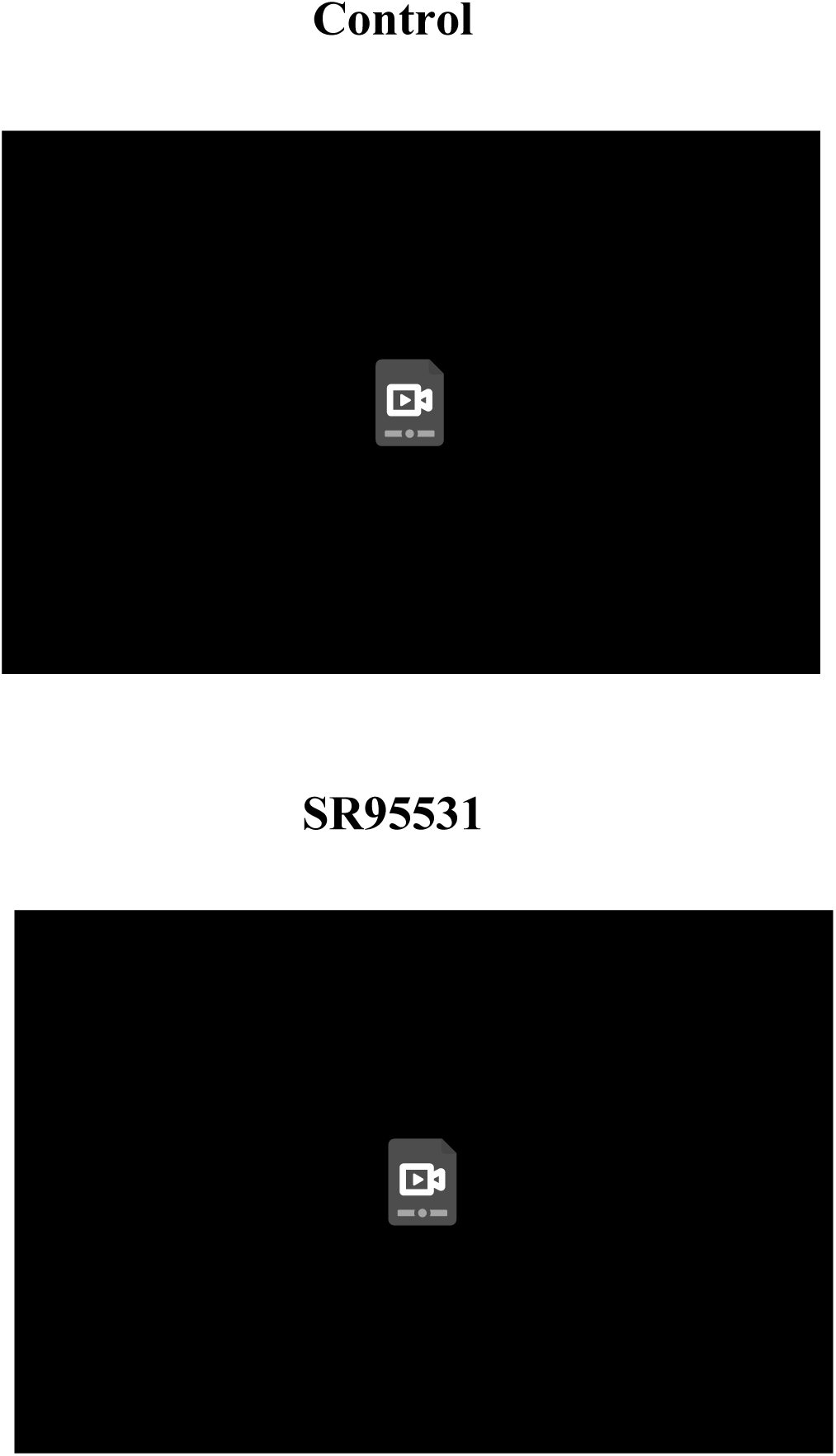
Time-lapse Ca²⁺ imaging of SACs in ChAT-Cre;Ai95 retinas before (control) and during SR95531. Corresponding heatmap videos are shown in Supplementary Videos 5–6.

**Supplementary Videos 5–6.**
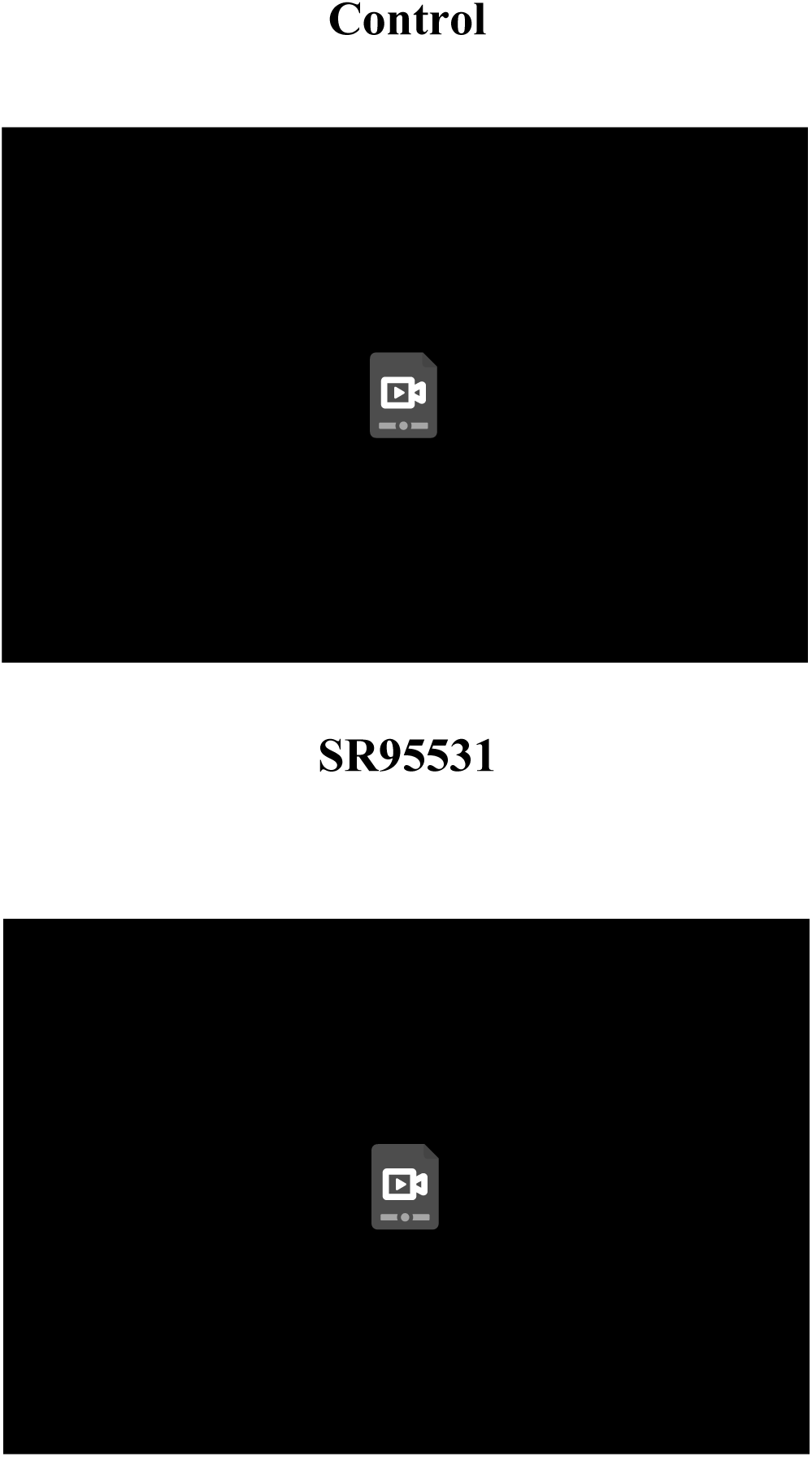
Corresponding heatmap videos for Fig. 6B show retinal waves before (control) and during SR95531 application, converted from time-lapse Ca²⁺ imaging videos in Supplementary Videos 3 and 4 using the MATLAB processing pipeline.

